# Mammalian TMC Family Proteins are Mechanically Gated Ion Channels

**DOI:** 10.64898/2026.08.18.745354

**Authors:** Songdi Fu, Jianying Dong, Xing Luo, Ting Xie, Wei Li, Yufei Luo, Zhiqiang Yan

**Author notes:** Correspondence: Zhiqiang Yan. These authors contributed equally to this work (Songdi Fu, Jianying Dong, Xing Luo).

## Abstract

Every known life form senses and reacts to mechanical forces. These mechanical stimuli can be converted into electrical signals by mechanically gated ion channels, a transduction cascade pivotal to numerous physiological functions including touch, hearing, mechanical pain, circulation, gastrointestinal function, and mechanical loading in various tissues. Despite continuous efforts, numerous mechanically gated ion channels with the mechanotransduction process underlying these physiological functions remain unidentified. Here, we focused on the transmembrane channel-like (TMC) protein family expressed in the cultured cells to identify those with potential mechanosensitive activity. Remarkably, in contrast to human TMC1/2 (HsTMC1/2), human TMC3-8 (HsTMC3-8) proteins are localized to the plasma membrane when heterologously expressed in the cultured cells. Further experiments revealed that mechanical poking stimuli can effectively activate HsTMC3-8. In addition, HsTMC3-8 induced stretch-activated currents and elicited well-resolved single-channel activities in response to negative pressure stimulation. The mutants near the putative pore region altered reversal potentials (Erev) of HsTMC3-8, suggesting that TMC3-8 are likely pore-forming subunits of ion channels. In summary, we proposed that TMC proteins are the largest mammalian mechanically gated ion channel family.

## Introduction

Cells detect and react to mechanical stimuli by converting them into electrical signals through a process called mechanotransduction, in mammals, mechanotransduction is critical for various processes including embryonic development, tactile perception, mechanical pain, proprioception, auditory function, regulation of blood flow, kidney flow sensing, lung development, maintenance of bone equilibrium, water homeostasis, as well as the progression of metastasis lesions^1–34^. Although the discovery of Piezo proteins in 2010^35^ marked a substantial breakthrough in the field, there exist mechanically gated ion channels remaining elusive, and their physiological functions are inadequately comprehended. This is particularly evident, as numerous sensations associated with both innocuous and noxious mechanical stimuli are independent of Piezo proteins in human and mouse^36–42^. Hence, identifying new mechanically gated ion channels is crucial.

Here we focused on members of the TMC protein family, which in mammals consists of eight members (TMC1-8)^43,44^. Among these, mammalian TMC1 and TMC2 have been identified as mechanically gated ion channels in heterologous expression systems and mediate mechanotransduction in hair cells^45,46^. In contrast, other TMC family members (TMC3-8) remain poorly characterized. Our results showed that, unlike HsTMC1/2, which are not localized in the plasma membrane of the cultured cells^47–52^, non-permeabilized staining and the biotinylated assay revealed that HsTMC3-8 are localized in the plasma membrane when heterologously expressed in Human Embryonic Kidney 293T (HEK293T) cells. Using poking stimuli, HsTMC3-8 produced robust mechanically activated (MA) current responses in the Piezo1-knockout (Piezo1-KO) HEK293T cells. Remarkably, when negative pressure pulses were applied in cell-attached mode, the expression of HsTMC3-8 could induce stretch-activated currents and exhibit clear single-channel activities in Piezo1-KO HEK293T cells. The reversal potentials of HsTMC3-8 mutants near the putative pore region shifted leftward compared to the WT HsTMC3-8 in Piezo1-KO HEK293T cells, suggesting that HsTMC3-8 are likely pore-forming subunits of ion channels. Previous studies suggested that mammalian TMC1/2 are mechanogated ion channels^45,46^. Taken together, these findings suggested that TMC proteins, comprising eight members (TMC1-8), represent the largest family of mechanically gated ion channels in mammals.

## Results

### Human TMC3-8 proteins are localized in the plasma membrane when heterologously expressed in the cultured cells

Mammalian TMC proteins family consists of eight members (TMC1-8) with highly conserved sequence alignments^43,44^ (Extended Data Fig. 1). As previously reported, mammalian TMC1/2 are not localized in the plasma membrane when heterologously expressed in the cultured cells^47–52^. Accordingly, whether the remaining mammalian TMC paralogs share similar localization characteristics remains unknown. To investigate their sub-cellular localization, we inserted an HA tag into the extracellular loop between TM1 and TM2 of HsTMC1-8, a position that we had uniquely identified as suitable in our previous study^45^. The HA tag was inserted after residues 236, 300, 189, 220, 489, 301, 226, and 166 of HsTMC1-8, respectively. Employing an anti-HA antibody and a membrane marker Wheat Germ Agglutinin (WGA) to perform non-permeabilized staining for HsTMC1-8, we found that, in contrast to HsTMC1/2, HsTMC3-8 proteins are localized in the plasma membrane of HEK293T cells (Fig. 1a). Notably, the signal was not uniformly distributed across the plasma membrane. Next, we assessed the plasma membrane expression of HsTMC1-8 using the cell-surface biotinylation assay. The results showed that HsTMC3-8 were expressed in the plasma membrane, whereas HsTMC1 and HsTMC2 were not, further supporting the conclusion that HsTMC3-8 are expressed in the plasma membrane of HEK293T cells (Fig. 1b). Notably, HsTMC3 exhibited detectable but less prominent plasma membrane localization compared with HsTMC4-8, possibly because HsTMC3 clusters with HsTMC1 and HsTMC2 in phylogenetic analyses^43,44^.

**Fig. 1.**
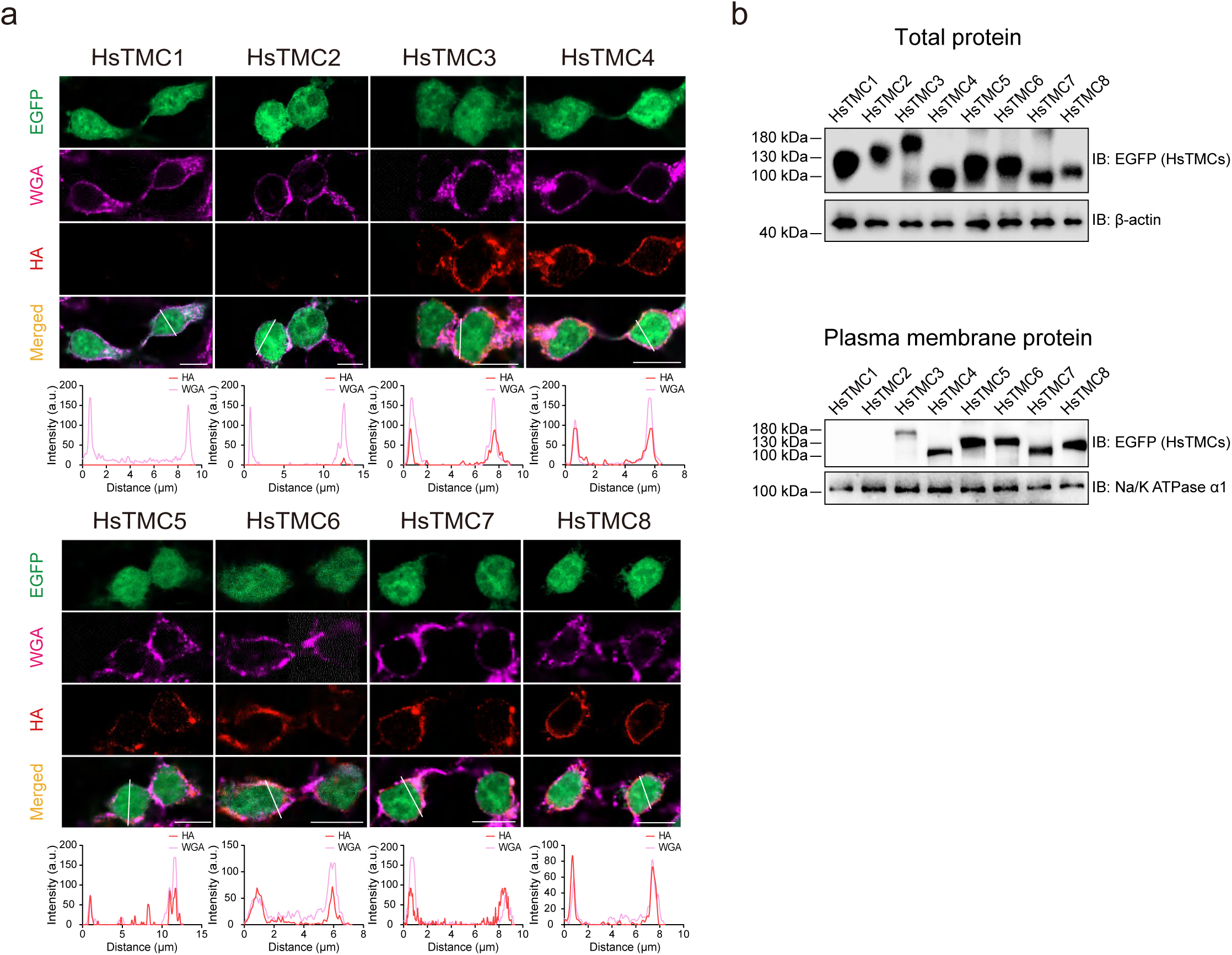
Human TMC3-8 localize in the plasma membrane in heterologous cells. a, Non-permeabilized staining of HsTMC1-8. HA tag is inserted in the extracellular loop between transmembrane helix 1 and 2 of HsTMC1-8. EGFP (green), expressed through an IRES vector, marks transfected cells. WGA (magenta) serves as a plasma membrane marker. HA (red) labels HsTMC proteins expressed in the plasma membrane. a.u., arbitrary units. Scale bars: 10 μm. **b,** Western blotting of EGFP-tagged HsTMC1-8 fusion proteins in biotinylated samples and total lysates using an anti-GFP antibody. β-actin served as a loading control for total proteins. Na/K ATPase α1, used as a loading control for membrane proteins, was detected on separate membranes processed in parallel with identical samples and experimental conditions. The experiment was independently repeated three times with similar results.

### Human TMC3-8 are mechanically gated ion channels

To test whether HsTMC3-8 are responsive to mechanical stimulation, we performed patch-clamp recordings by expressing HsTMC3-8 in the mechanically insensitive Piezo1-KO HEK293T cells^53,54^. Notably, HsTMC3-8 proteins were activated by mechanical stimulation with an 8 μm poking displacement at a holding potential of -60 mV, showing positive ratios of 45.45%, 50.00%, 46.15%, 55.56%, 50.00%, and 53.85%, respectively (Fig. 2a,b). This indicated that TMC3-8 may function as mechanogated ion channels. To investigate the pharmacological properties of HsTMC3-8, we opted for five ion channel blockers^1,55–66^: gadolinium chloride (GdCl_3_) is known for its ability to inhibit cation channels; ruthenium red (RR) is a universal pore blocker for a wide array of cation channels; dihydrostreptomycin (DHS) is a permeant blocker for the mechanotransduction channels in hair cells; GsMTx-4 is a specific blocker of mechanosensitive ion channels; and 5-nitro-2-(3-phenylpropylamino) benzoic acid (NPPB) is a blocker of Cl^-^ channels. The currents associated with HsTMC3-8 in Piezo1-KO HEK293T cells were effectively inhibited by GdCl_3_, RR, DHS, and GsMTx-4, whereas NPPB had no effect (Fig. 2c). When investigating the ion selectivity of HsTMC3-8, MA currents were attenuated in the presence of N-methyl-D-glucamine chloride (NMDG-Cl) solution but remained unaffected in Na-gluconate solution (Fig. 3a-f), suggesting that HsTMC3-8 are mechanosensitive cation channels.

**Fig. 2.**
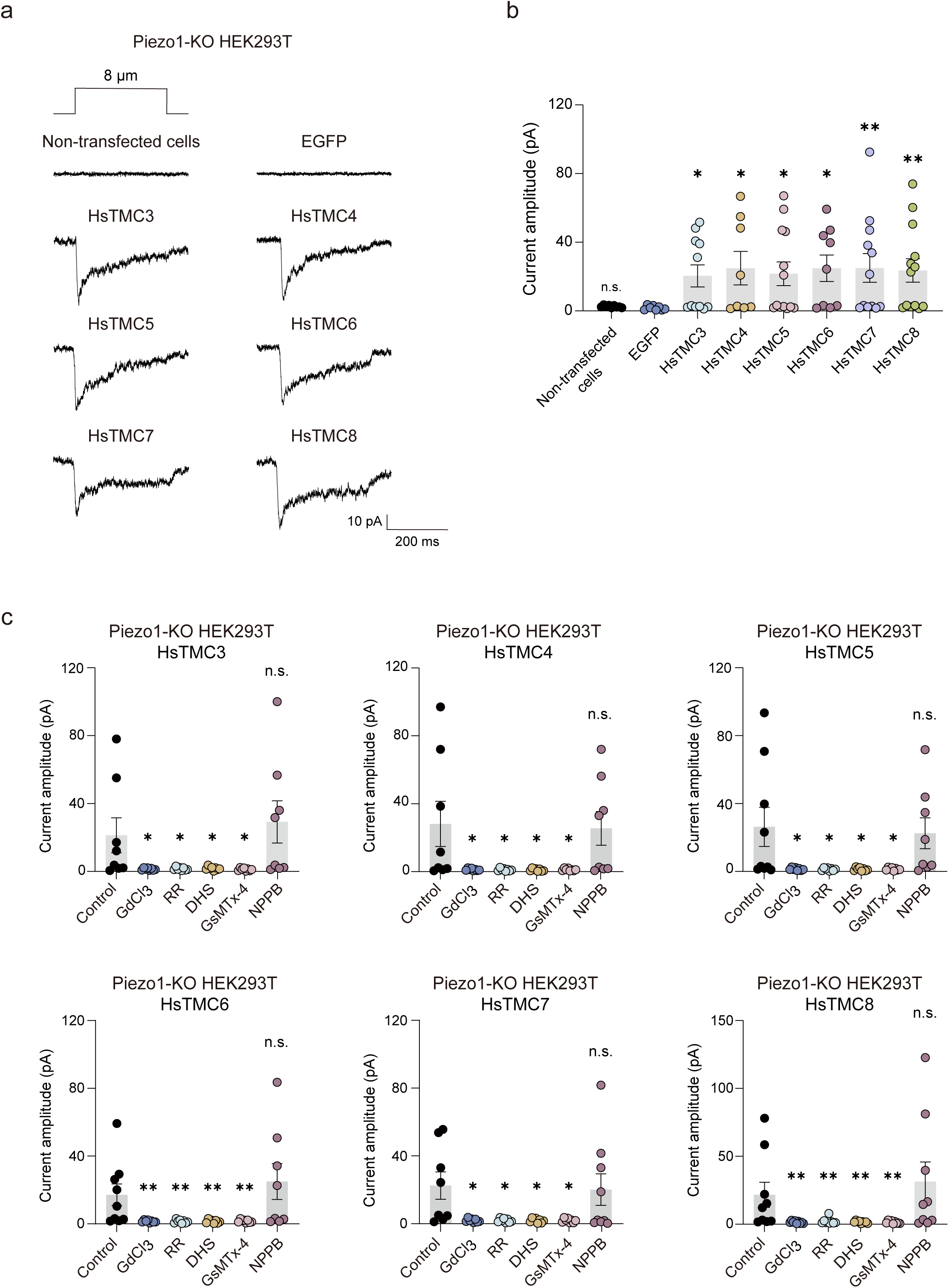
Human TMC3-8 can be activated by poking stimuli. a, Representative current traces of the non-transfected cells, EGFP, and HsTMC3-8 in response to 8 μm displacement at a holding potential of -60 mV in Piezo1-KO HEK293T cells. **b,** Current amplitudes of non-transfected cells, EGFP, and HsTMC3-8. n ≥ 8, n represents the number of cells. All error bars: mean ± SEM. All comparisons were made against the EGFP group. \*\**p* < 0.01, \**p* < 0.05, *p* ≥ 0.05, n.s., not significant, Mann-Whitney test, unpaired *t* test, or unpaired *t* test with Welch’s correction. **c,** Human TMC3-8 mechanically activated currents evoked by 8 μm poking stimulation were inhibited by GdCl₃, RR, DHS, and GsMTx-4, but not by NPPB, in Piezo1-KO HEK293T cells. n ≥ 8, n represents the number of cells. \*\**p* < 0.01, \**p* < 0.05, *p* ≥ 0.05, n.s., not significant, Mann-Whitney test or unpaired *t* test with Welch’s correction. Data for control and blocker-treated conditions were collected independently using different cells. All error bars: mean ± SEM.

**Fig. 3.**
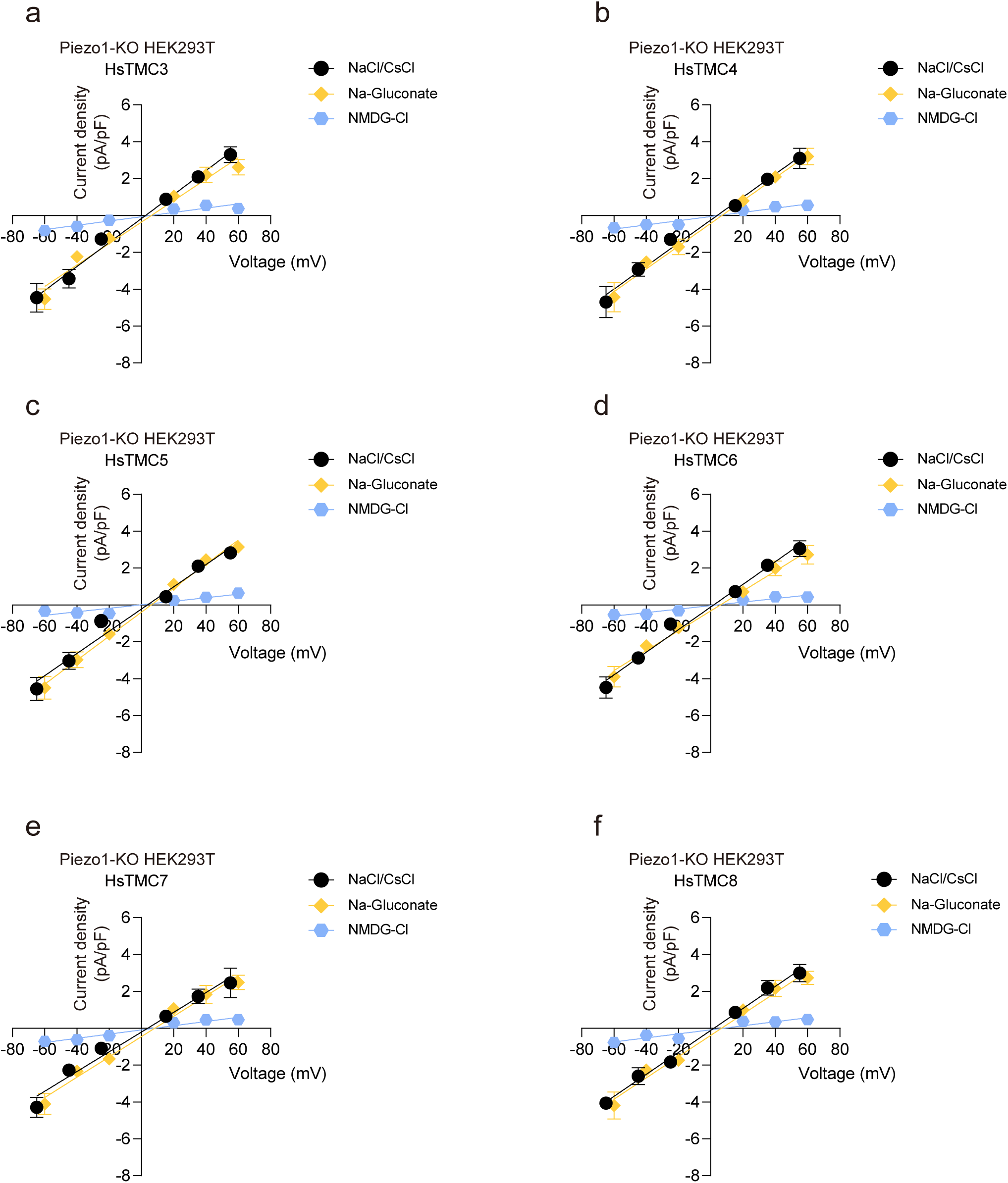
Human TMC3-8 function as mechanosensitive cation channels. a-f, The current density-voltage relationships of HsTMC3-8 evoked by mechanical stimuli in response to an 8 μm displacement from -60 mV to +60 mV. The currents were recorded in NaCl/CsCl, Na-gluconate, and NMDG-Cl solutions, respectively. MA currents were recorded at each voltage. n ≥ 9, n represents the number of cells recorded at each voltage. All error bars: mean ± SEM. The liquid junction potentials have been subtracted from the current density-voltage relationship statistics.

### Human TMC3-8 show stretch-activated currents and single-channel activities

After assessing mechanical sensitivity utilizing whole-cell recording, we further examined the HsTMC3-8-induced currents in Piezo1-KO HEK293T cells using an alternative mechanosensitivity assay, which applied negative pressure pulses through the recording pipette in the cell-attached mode. Heterologous expression of HsTMC3-8 in Piezo1-KO HEK293T cells induced progressively increasing stretch-activated currents with increasing pressure stimulation at -60 mV (Fig. 4a), while no pressure-induced currents were observed in non-transfected or EGFP-transfected Piezo1-KO HEK293T cells (Fig. 4b). The half-maximal activation pressures (P_50_) of HsTMC3-8 were -87.09 ± 4.51 mmHg, -86.56 ± 6.08 mmHg, -92.01 ± 6.31 mmHg, -88.98 ± 7.01 mmHg, -87.84 ± 6.66 mmHg and -80.81 ± 9.78 mmHg (Fig. 4a and 4c), respectively. Furthermore, HsTMC3-8 exhibited a significant increase in single-channel activities (Fig. 5a), while no single-channel activities were observed in non-transfected or EGFP-transfected Piezo1-KO HEK293T cells (Fig. 5b). The clear single-channel activities were well revealed, showing the conductance of HsTMC3-8 as follows: 41.21 ± 1.82 pS, 35.23 ± 1.23 pS, 36.81 ± 1.89 pS, 42.18 ± 2.76 pS, 38.70 ± 1.46 pS, and 38.87 ± 0.92 pS, respectively. Notably, the single-channel open probabilities of HsTMC3-8 significantly increased in response to applied pressure (Fig. 5c). Taken together, these findings indicated that HsTMC3-8 ion channels can be activated by mechanical forces, consistent with the results observed from poking stimulation in the whole-cell recording mode.

**Fig. 4.**
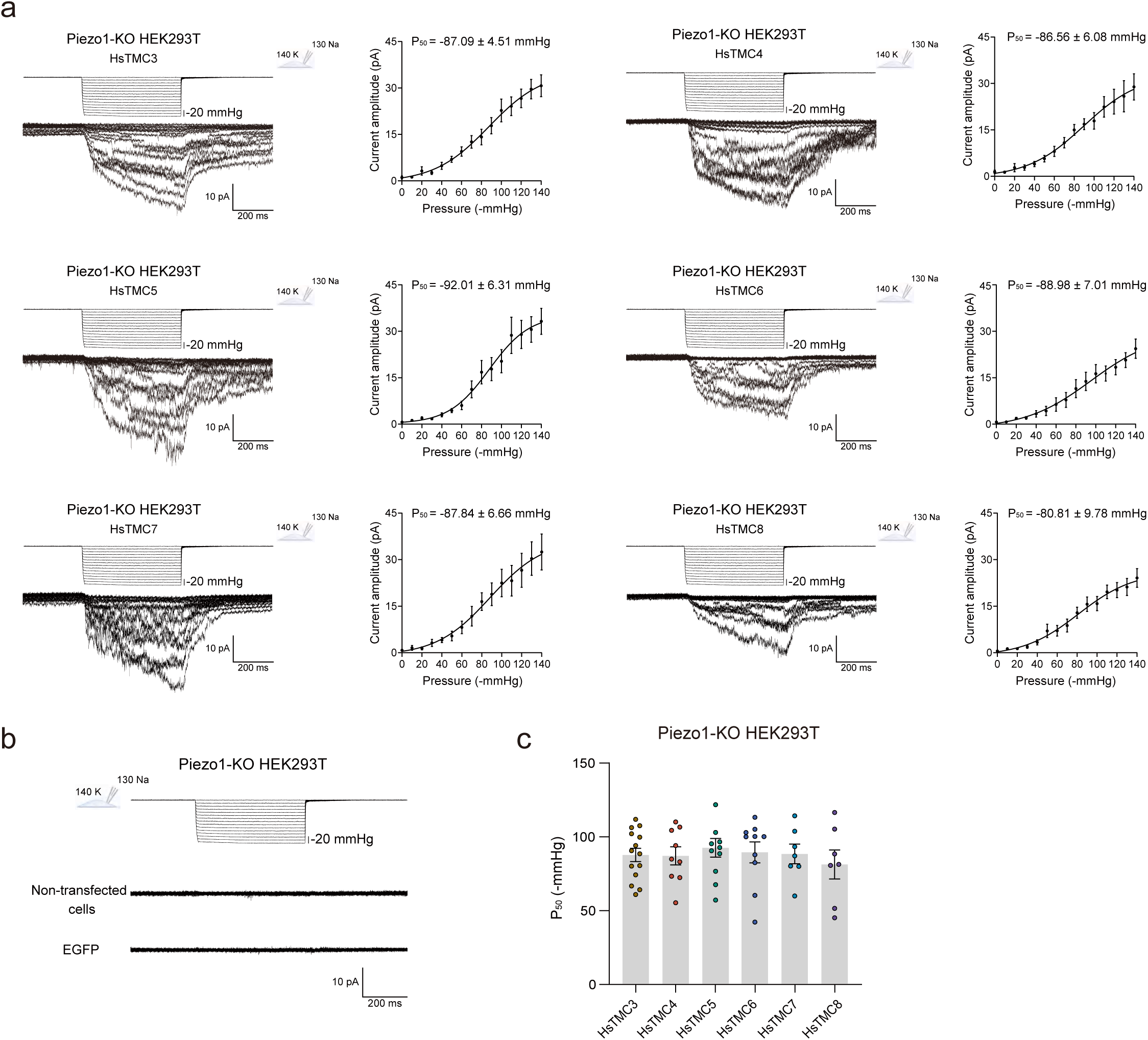
Human TMC3-8 could be activated by stretch stimuli. a, Left: Representative traces of stretch-activated currents in the cell-attached recording mode from Piezo1-KO HEK293T cells transfected with human TMC3-8. Right: Pressure-response current curves fitted with the Boltzmann equation. n ≥ 7, n represents the number of cells. Negative pressures were applied from 0 mmHg to -140 mmHg with -10 mmHg per step. The patch membrane was held at -60 mV. All error bars: mean ± SEM. **b,** Representative traces of stretch-activated currents in the cell-attached mode of non-transfected and EGFP-transfected Piezo1-KO HEK293T cells. **c,** P_50_ values obtained from human TMC3-8 in Piezo1-KO HEK293T cells. n ≥ 7, n represents the number of cells. All error bars: mean ± SEM.

**Fig. 5.**
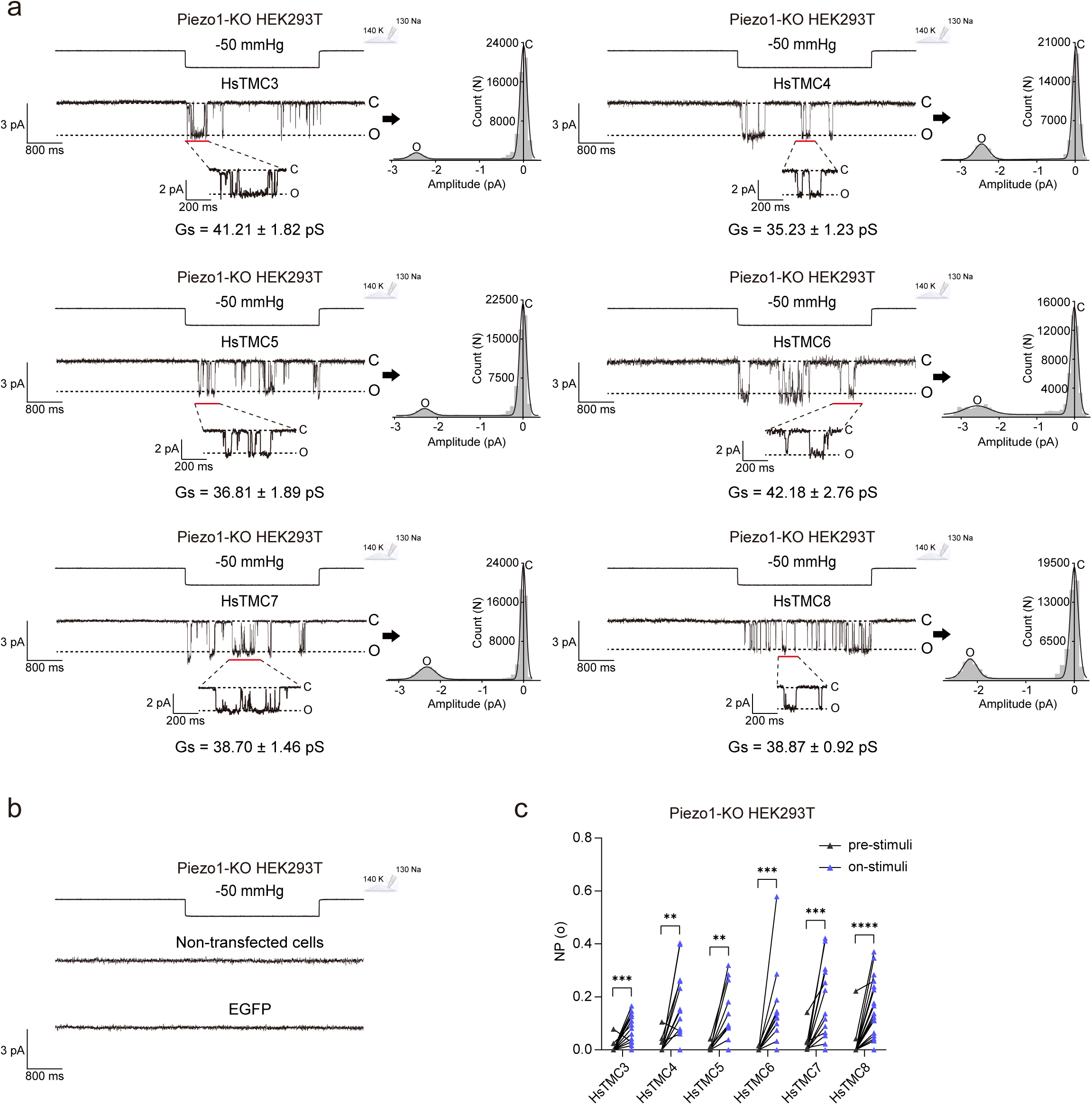
Human TMC3-8 exhibit negative pressure-induced single-channel activities. a,b, Representative single-channel current traces of human TMC3-8, non-transfected and EGFP-expressed in Piezo1-KO HEK293T cells at -60 mV under the -50 mmHg negative pressure for 3 s. C, closed; O, open. The traces marked by red outlines are enlarged and displayed at the bottom. All-point current histograms of the single-channel recordings acquired under -50 mmHg pressure for 3 s. **c,** Single-channel open probabilities of human TMC3-8 at -60 mV with pressure stimulation. n ≥ 18, n represents the number of cells. \*\*\*\**p* < 0.0001, \*\*\**p* < 0.001, \*\**p* < 0.01, Wilcoxon matched-pairs signed rank test.

### Human TMC3-8 are pore-forming subunits of mechanically gated ion channels

To test whether human TMC3-8 function as pore-forming subunits of ion channels, we generated point mutants by substituting glutamic acid (Glu, E) with alanine (Ala, A) in HsTMC3-8: HsTMC3-E501A, HsTMC4-E477A, HsTMC5-E726A, HsTMC6-E542A, HsTMC7-E488A, and HsTMC8-E420A. These substitutions were based on a highly conserved cysteine (Cys, C)-tryptophan (Trp, W)-Glu (CWE) motif from TMC signature sequence^43^ and are hypothesized to be located near the predicted pore region in the TM6 domain of TMC proteins^43,67–70^ (Fig. 6a and Extended Data Fig. 1). In addition, we selected key amino acid residues in the pore region based on a study that identified a series of functionally important sites in TMC1^70^. Through sequence alignment, we found that the conserved sites corresponding to the amino acids in HsTMC3-8: HsTMC3-D515A, HsTMC4-D491A, HsTMC5-D740A, HsTMC6-D556A, HsTMC7-D502A, and HsTMC8-T438A (Fig. 6a). Remarkably, the current density-voltage relationships of wild-type and point-mutated HsTMC3-8 MA currents in Piezo1-KO HEK293T cells exhibited linearity within the range of -60 mV to +60 mV, with reversal potentials of HsTMC3-8 mutants shifted leftward compared to the wild-type HsTMC3-8: HsTMC3, from 2.30 mV to -1.09 mV (E501A) and 1.47 mV (D515A); HsTMC4, from 2.88 mV to -0.21 mV (E477A) and 1.33 mV (D491A); HsTMC5, from 2.91 mV to -2.26 mV (E726A) and 1.42 mV (D740A); HsTMC6, from 2.73 mV to - 1.23 mV (E542A) and -3.11 mV (D556A); HsTMC7, from 3.24 mV to -2.83 mV (E488A) and 0.62 mV (D502A); HsTMC8, from 2.62 mV to -1.99 mV (E420A) and 1.22 mV (T438A) (Fig. 6b-g). These results indicated that human TMC3-8 are likely the pore-forming subunits of ion channels that can be activated by mechanical stimuli.

**Fig. 6.**
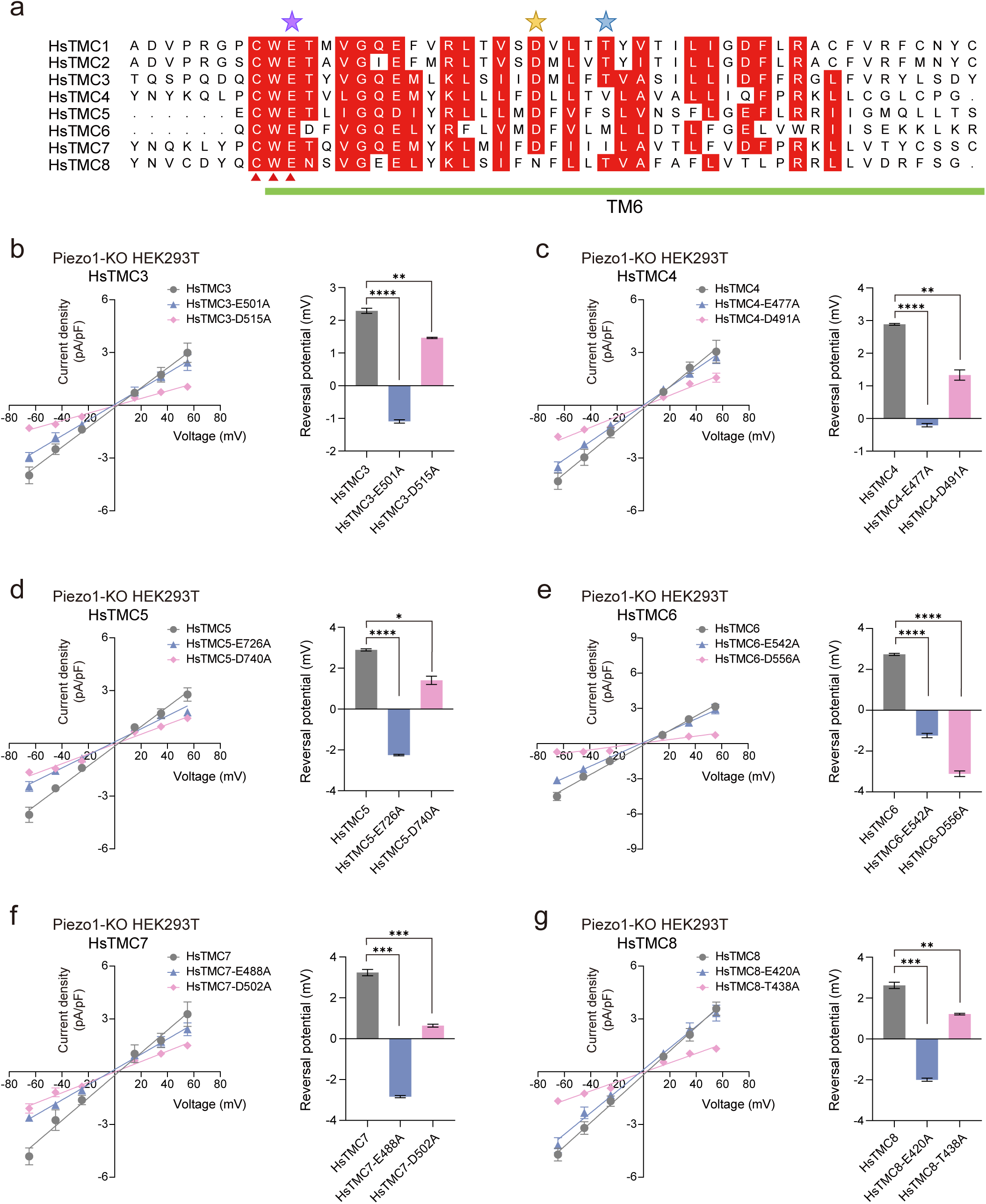
Human TMC3-8 are likely pore-forming subunits of mechanically gated cation channels in Piezo1-KO HEK293T cells. a, Amino acid sequence alignment of the TM6 region in human TMC1-8 proteins (HsTMC1: NP_619636.2; HsTMC2: NP_542789.2; HsTMC3: NP_001074001.1; HsTMC4: NP_653287.2; HsTMC5: NP_001248770.1; HsTMC6: NP_009198.4; HsTMC7: NP_079123.3; HsTMC8: NP_689681.2). Identical and highly conserved amino acid residues were marked in red boxes. The CWE motif from TMC signature sequence was indicated with red triangles. The TM6 region was denoted with a green bar. Purple star indicates the point mutation sites in the highly conserved CWE motif of HsTMC3-8; yellow star indicates point mutation sites for HsTMC3-7; blue star indicates the point mutation site for HsTMC8. Sequence alignment was generated in ESPript 3.0 and Clustal Omega. **b-g,** Left: The current density-voltage relationships of wild-type and point-mutated HsTMC3-8 in Piezo1-KO HEK293T cells evoked by mechanical stimuli in response to an 8 μm displacement from -60 mV to +60 mV, respectively. MA currents were recorded at each voltage. n ≥ 9, n represents the number of cells recorded at each voltage. Junction potentials were calculated based on the solution composition and corrected before plotting. Right: the mutants shifted the reversal potential of wild-type HsTMC3-8. All error bars: mean ± SEM. n = 3, n represents three independent experiments. \*\*\*\**p* < 0.0001, \*\*\**p* < 0.001, \*\**p* < 0.01, \**p* < 0.05, unpaired *t* test with Welch’s correction.

## Discussion

In our quest to identify new mechanically gated ion channels, coupled with our ongoing interest in the TMC protein family, we conducted non-permeabilized staining and the biotinylated assay of TMC3-8 proteins in HEK293T cells, revealing an intriguing distinction, unlike TMC1/2^47–52^, TMC3-8 are effectively localized to the plasma membrane when heterologously expressed in the cultured cells. Patch-clamp recordings revealed that human TMC3-8 proteins from the TMC family exhibited mechanically activated currents in response to poking stimuli in Piezo1-KO HEK293T cells. Notably, when applying negative pressure pulses in the cell-attached mode, HsTMC3-8 expressed in Piezo1-KO HEK293T cells showed stretch-activated currents and clear single-channel activities. More importantly, the reversal potentials of HsTMC3-8 mutants near the putative pore region shifted leftward compared to the wild-type HsTMC3-8 in Piezo1-KO HEK293T cells, suggesting that HsTMC3-8 are likely pore-forming subunits of ion channels.

The identification of mechanically gated ion channels in TMC3-8 marks the beginning of a significant exploration into the physiological roles of TMC family proteins. The seminal discovery that Piezo1/2 are mechanically gated ion channels^35^ has sparked extensive investigations into the physiological functions of Piezo1/2. These comprise their involvements in various aspects such as lung inflation^71,72^, vascular structure^73,74^, curvature and lateral tension in the blood cells^75^, epithelial cells division and extrusion^76^, touch^77–79^, and itch^80^. Mammalian TMC proteins family has eight members (TMC1-8) with high homology^43,44^, at present, the roles of mammalian TMC family members in various physiological functions are progressively being discovered. TMC1/2, found at the stereocilia tips of the hair cells, are proposed as the mechanotransduction channels in hair cells in the vertebrate auditory system^81–104^, their invertebrate homologs also play vital roles in various physiological functions^51,68,69,105–112^. Lung-innervating TMC3-positive neurons can induce broncho-constriction and dilation in the respiratory cycle^113^. Microvilli in Intestinal Epithelial Cells (IECs) are interconnected via extracellular cadherin-based tip-links similar to stereocilia, and act as mechanosensors. TMC4 and TMC5 proteins are potential microvillar-mechanosensitive ion channels, located along with Calcium and Integrin Binding Protein 3 (CIB3) at the distal end of microvilli^114^. TMC6 or TMC8 homozygous mutations induce a severe genetic skin disorder called Epidermodysplasia Verruciformis (EV), in which half of the cases develop into skin carcinomas, due to a specific susceptibility to related human papillomaviruses^44,115–118^. TMC7 mutant mice exhibited male infertility and severe defects in spermiogenesis^119,120^. A study also suggested that TMC3, TMC5, and TMC7 are associated with nociception, whereas TMC5 and TMC7 additionally play roles in itch^121^. Hence, following the discovery that the TMC family proteins function as mechanically gated ion channels, their broader physiological functions and the relationship between these functions and the mechanosensitivity of TMC proteins should be explored in depth.

In summary, we identified mammalian TMC3-8 as robust mechanically gated ion channels by electrophysiological recording, revealing the inherent properties of the TMC family proteins. Our discoveries might open avenues for investigating the gating mechanisms of TMC3-8 and their interactions with other components of the mechanoelectrical transduction pathway for translating mechanical stimuli and pave the way to explore the functions of TMC3-8 across various physiological processes.

### Methods Cell lines

HEK293T cells and Piezo1-KO HEK293T cells (from our paper^45^, RRID: CVCL_D4Z8), were cultured in Dulbecco’s Modified Eagle Medium (DMEM) supplemented with 10% fetal bovine serum (FBS), and 1% Penicillin/Streptomycin (P/S) at 37℃ with 5% CO_2_. The cells were plated onto 10 mm round glass coverslips in 24-well plates before transfection. Transfection was carried out according to the manufacturer’s instructions (Lipofectamine 3000) after 12-24 h. Genes were synthesized or obtained from the gene libraries (CCSB-Broad Lentiviral Expression Library, Precision LentiORF whole genome expression library, and Human hORFeome V8.1 Library, Horizon).

### Molecular cloning

For non-permeabilized staining and patch-clamp recording experiments, all genes were synthesized and subcloned into the pIRES2 vector, with EGFP introduced at the C-terminus for non-fusion expression. For non-permeabilized staining, an HA tag was inserted into the extracellular loop between TM1 and TM2 of each TMC protein, after residues 236, 300, 189, 220, 489, 301, 226, and 166 of HsTMC1-8, respectively. For western blot analyses, HsTMC genes were cloned into the pN3 vector to generate N-terminal EGFP fusion constructs. All cDNAs were verified by sequencing for the full length.

### Non-permeabilized staining for HEK293T cells

HEK293T cells underwent transfection approximately 24-48 h prior to immunostaining. Cells were labeled with WGA for 10 min at 37°C. After labeling, the solution was removed, and the cells were washed three times with 1× PBS. The cells were then fixed with a 4% Paraformaldehyde (PFA) solution for 15 min at room temperature (25℃) and subsequently blocked with 2% Bovine Serum Albumin (BSA) for 1 h at room temperature. The cells were then incubated with the primary antibody (anti-HA mouse monoclonal antibody, 1:1000) overnight at 4℃. Detection of the primary antibody was accomplished utilizing the secondary antibody (goat anti-mouse IgG, 1:1000) for 1 h at room temperature (25℃). Hoechst (1:400) was then incubated for 12 min at room temperature. Images were captured utilizing a ZEISS LSM 980 confocal microscope. The mean fluorescence intensity of surface-expressed HsTMC proteins was quantified using ImageJ by threshold-based segmentation of membrane regions, followed by automatic identification and measurement of fluorescence signals.

### Protein extraction

For whole-cell protein extraction, sample lysates were extracted using Radio Immunoprecipitation Assay (RIPA) lysis buffer with 1× protease inhibitor cocktail and 1 mM Phenylmethylsulfonyl Fluoride (PMSF). After the cells were vortexed gently, the cell debris was removed by centrifugation at 15,000 rpm for 6 min at 4°C.

For plasma membrane protein extraction, sample lysates were prepared using a Pierce^TM^ Cell Surface Protein Biotinylation and Isolation Kit. The kit utilizes Sulfo-NHS-SS-Biotin to covalently label primary amines on the cell surface. This membrane-impermeable reagent contains a cleavable disulfide bond that enables mild elution under reducing conditions using Dithiothreitol (DTT). Biotinylated proteins are enriched through high-affinity binding to NeutrAvidin agarose resin and subsequently eluted for downstream analyses. Briefly, cells cultured in 15 cm dishes were first washed with PBS and then incubated with 10 mL of EZ-Link™ Sulfo-NHS-SS-Biotin solution at room temperature for 10 min to biotinylate primary amines exposed on the cell surface. To terminate the biotinylation reaction, cells were extensively washed with ice-cold TBS to remove unreacted Sulfo-NHS-SS-Biotin, followed by centrifugation at 500 ×g for 3 min at 4°C. Cell pellet was subsequently lysed with 500 µL of lysis buffer containing protease inhibitors for 45 min and then centrifuged cell lysates at 15,000 ×g for 5 min at 4°C, and the clarified supernatant was transferred to a new 1.5 mL tube.

The clarified supernatant was used to capture the labeled protein with 250 µL NeutrAvidin agarose. Subsequently, mix 225 µL Elution Buffer and 25 µL DTT stock solution and add 200 µL to the resin and cap the column to elute proteins. Then incubate the mixture for 30 min at room temperature with end-over-end mixing on a rotator. Then loosen the top cap, remove the bottom cap, place into a collection tube and centrifuge column for 2 min at 1,000 ×g. The eluted products were further enriched using 15 μL of magnetic beads-conjugated anti-GFP VHH single domain antibody and incubated at 4℃ with rotation for at least 8 h. Then the beads were washed three times with wash buffer (50 mM Tris-HCl, 150 mM NaCl, pH 7.5) and resuspended in 30 μL of 2× SDS loading buffer for subsequent western blot analysis.

### Western blot analysis

For western blot analysis, protein samples were separated by 12.5% SDS-PAGE and transferred to a Nitrocellulose (NC) membrane or Polyvinylidene Fluoride (PVDF) membrane. The membranes were blocked with 5% non-fat powdered milk in TBST and incubated with the primary antibodies diluted in dilution buffer at 4℃ overnight. After washing the membranes with TBST for three times, HRP-conjugated secondary antibody was added and incubated together for 1 h at room temperature and then washed again with TBST. The antibody-antigen complex was visualized using an enhanced chemiluminescent kit.

The primary antibodies included rabbit anti-GFP (1:900), rabbit anti-β-actin (1:4000), and rabbit anti-Na/K ATPase α1 (1:900). The secondary antibody was HRP-conjugated goat anti-Rabbit IgG (H+L) (1:7000). β-actin and Na/K ATPase α1 were used as loading controls. The results were representative of three individual experiments in triplicate.

### Patch-clamp recording

Patch-clamp recordings were performed in Piezo1-KO HEK293T cells 24 h after transfection. For whole-cell recordings, poking stimuli were transmitted to the cell utilizing a stimulation pipette driven by the Piezo Controller. The stimulation pipette was sealed and fire-polished to achieve a tip diameter of approximately 2 μm. Currents were sampled at 10 kHz and filtered at 3 kHz. The pipette buffer was composed of (in mM): 133 CsCl, 1 MgCl_2_, 1 CaCl_2_, 5 EGTA, 4 MgATP, and 10 HEPES (300-310 mOsm, pH 7.3 adjusted with CsOH). The bath buffer consisted of (in mM): 133 NaCl, 2.5 CaCl_2_, 3 KCl, 1 MgCl_2_, 15 D-glucose, 10 HEPES (300-310 mOsm, pH 7.3 adjusted with NaOH). Mechanically activated currents were evoked using an 8 μm poking stimulus lasting 300 ms at a holding potential of -60 mV. For recording in the NMDG-Cl or Na-gluconate solution, equimolar N-methyl-D-glucamine chloride (NMDG-Cl) and Na-gluconate, respectively, was used to replace NaCl in the bath solution and replace CsCl in the pipette solution. For blocker experiments, 10 μM GdCl_3_, 30 μM RR, 200 μM DHS, 5 μM GsMTx-4, and 100 μM NPPB were tested. Data for control and blocker-treated conditions were collected independently using different cells. The current density-voltage curves were fitted over the range of -60 mV to +60 mV. Throughout this procedure, each cell was subjected to one stimulation at each voltage, and liquid junction potentials were calculated using Clampex 11.2.

For cell-attached recordings, membrane patches were stimulated with negative pressure pulses applied through the recording electrode utilizing an HSPC-1 device. Currents were sampled at 20 kHz and filtered at 1 kHz. Pipettes were filled with a solution containing (in mM): 130 NaCl, 5 KCl, 10 HEPES, 1 CaCl_2_, 1 MgCl_2_, and 10 Tetraethylammonium chloride (TEA-Cl), with the pH adjusted to 7.3 utilizing NaOH. The external solution contained (in mM) 140 KCl, 10 HEPES, 1 MgCl_2_, and 10 D-glucose, with the pH adjusted to 7.3 utilizing KOH. For pressure-step recordings, negative pressure was applied through the recording electrode as 500-ms steps ranging from 0 to -140 mmHg in 10-mmHg increments. Single-channel current amplitude characterization was performed on cells held at -60 mV, with a negative pressure of -50 mmHg applied for 3 s.

For all patch-clamp experiments, recording pipettes with a resistance range of 3 to 5 MΩ were used. Seal resistance was measured between 1 and 2 GΩ. All experiments were conducted at a constant temperature of approximately 25℃.

## Acknowledgments

We thank Bioimaging Core of Shenzhen Bay Laboratory for providing imaging support. **Fundings:** This work is supported by funds from STI2030-Major Projects (2021ZD0203304); Shenzhen Medical Research Funding (B2302032); National Natural Science Foundation of China (32525033, 31970931); the National Key R&D Program of China Project (2021YFA1101302); and Shenzhen Science and Technology Program (RCJC20210609104631084).

## Author contributions

Z.Y. conceived the project and designed the experiments. S.F. and Z.Y. wrote the manuscript. S.F. performed the non-permeabilized staining; S.F. and X.L. performed biotinylation assays; S.F., J.D., X.L., T.X., W.L., and Y.L. performed the patch-clamp recording experiments. All authors discussed the results and commented on the manuscript.

## Data availability

This paper does not report original code. Any additional information required to reanalyze the data reported in this paper is available from the lead contact upon request.

## Notes

### Competing Interest Statement

The authors have declared no competing interest.

