## Extended Data Fig. 1 | for "Mammalian TMC Family Proteins are Mechanically Gated Ion Channels"

**This PDF file includes:**

**Extended Data Fig. 1 | Amino acid sequence of human TMC1-8.**

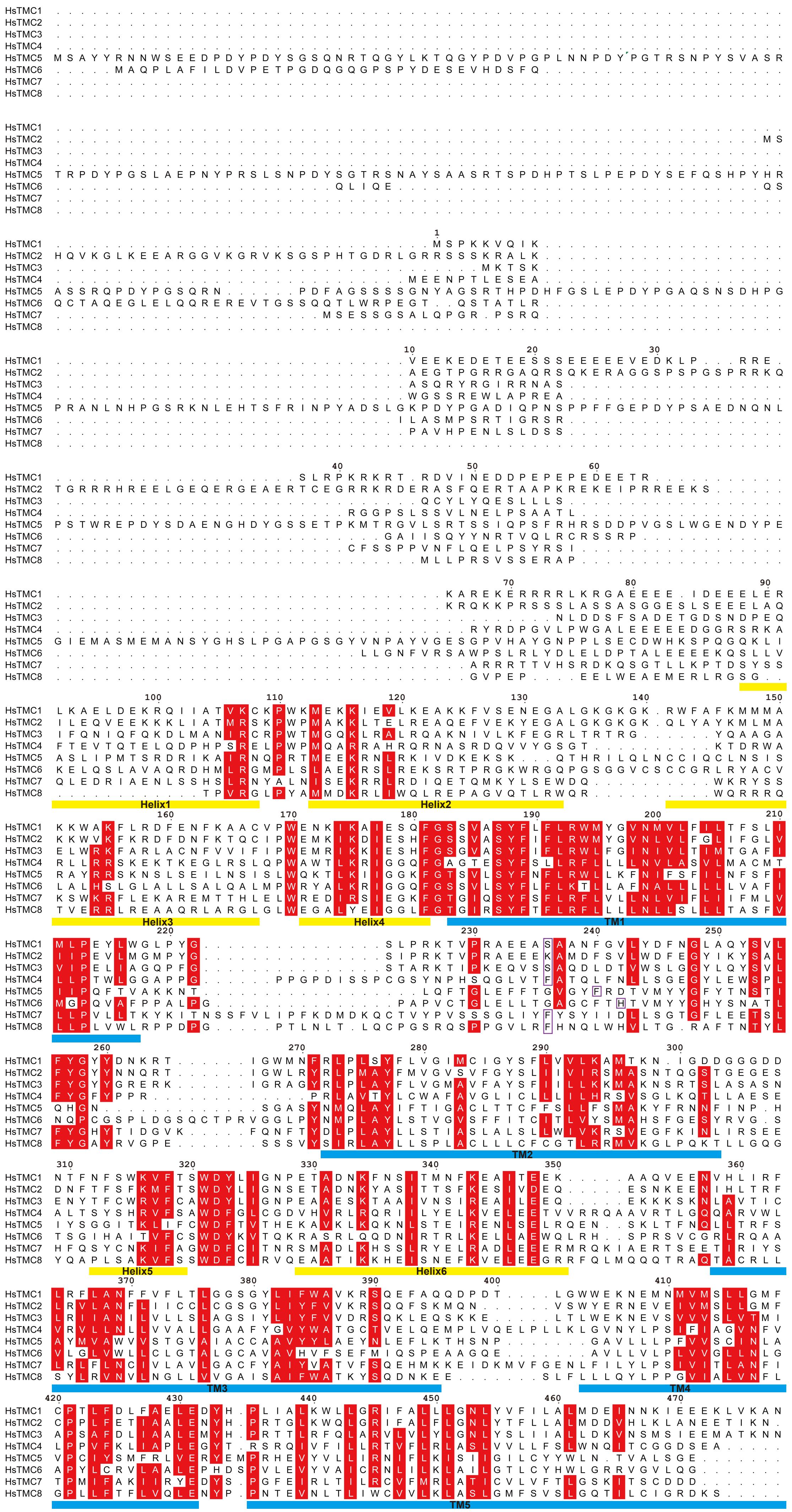


Continued on next page

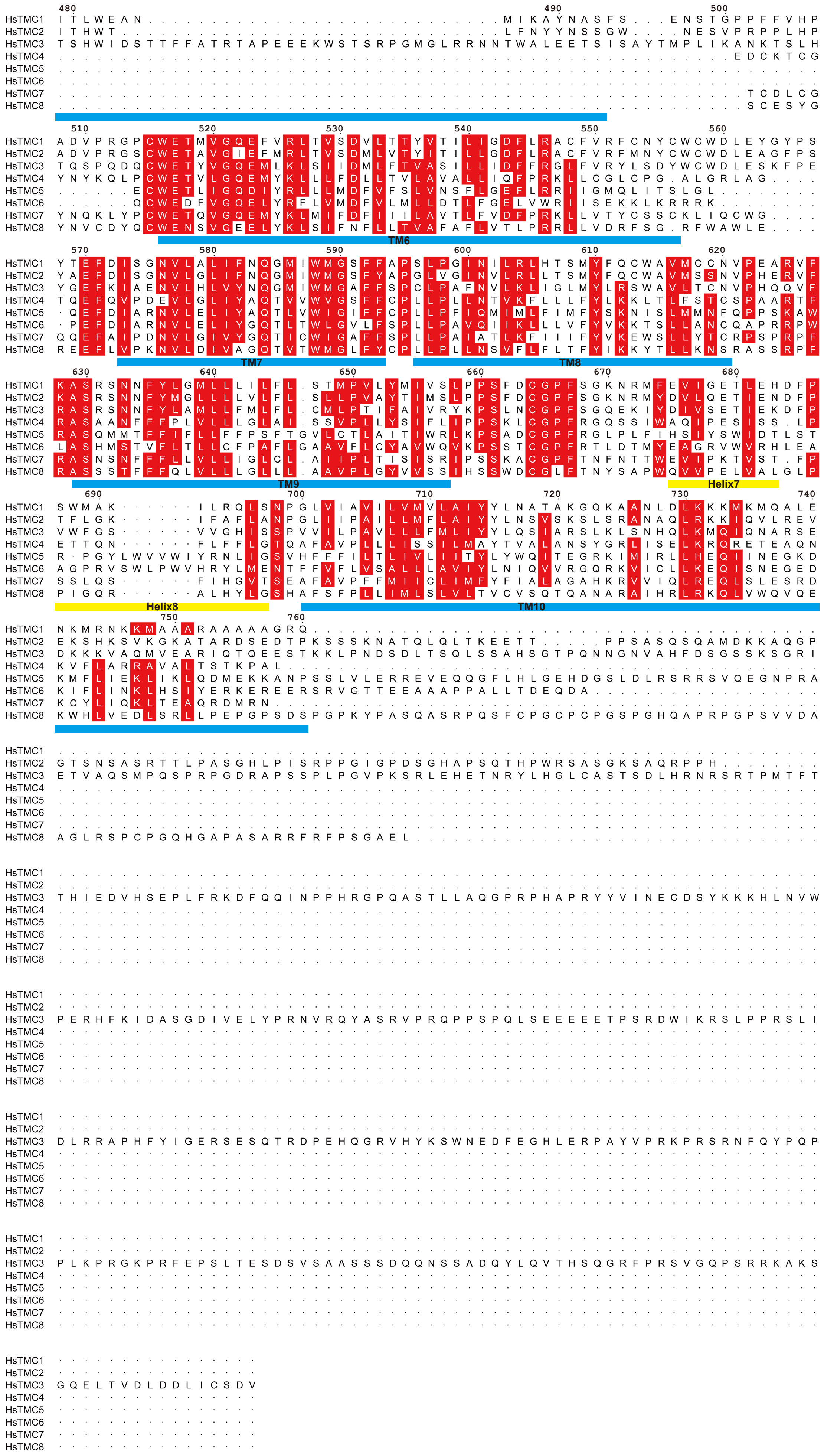


Extended Data Fig. 1

**Extended Data Fig. 1 | Amino acid sequence of human TMC1-8.** Amino acid sequence alignment of the human TMC1-8 proteins (HsTMC1: NP_619636.2; HsTMC2: NP_542789.2; HsTMC3: NP_001074001.1; HsTMC4: NP_653287.2; HsTMC5: NP_001248770.1; HsTMC6: NP_009198.4; HsTMC7: NP_079123.3; HsTMC8: NP_689681.2). Putative transmembrane domains and helices were based on the alignment with *Caenorhabditis elegans* TMC1 (NP_508221.3), which has a resolved structure (PDB-CPX-114691). Identical and highly conserved amino acid residues were marked in red boxes. The transmembrane regions were marked with blue bars and helices were marked with yellow bars. For non-permeabilized immunofluorescence staining, the HA tags were inserted after the amino acid residues indicated by purple boxes. The marked domains are based on the amino acid sequence of HsTMC1. Sequence alignment was generated in ESPript 3.0 and Clustal Omega.
